# fishROI: A specialized workflow for semi-automated muscle morphometry analysis in teleosts

**DOI:** 10.64898/2026.03.27.714781

**Authors:** Yansong Lu, Michael Pan, Vijayishwer Jamwal, Jake Locop, Avnika Ruparelia, Peter Currie

**Affiliations:** Australian Regenerative Medicine Institute, Monash University, Wellington Road, Clayton, VIC 3800, Australia; Melbourne Veterinary School, The University of Melbourne, Corner Flemington Road and Park Drive, Parkville, VIC 3052, Australia; MDI Biological Laboratories, Old Bar Harbor Rd, Bar Harbor, ME 04609, United States; Department of Anatomy and Physiology, School of Biomedical Sciences, Faculty of Medicine Dentistry and Health Sciences, University of Melbourne, Melbourne, VIC 3010, Australia; Centre for Muscle Research, Department of Anatomy and Physiology, University of Melbourne, Melbourne, Victoria, 3010, Australia; EMBL Australia, Victorian Node, Level 1, 15 Innovation Walk, Monash University, Wellington Road, Clayton, VIC 3800, Australia

## Abstract

Quantitative histological analysis of skeletal muscle morphometry provides critical insights into muscle physiology but remains labor-intensive and technically demanding. While recent developments in machine-learning-based image segmentation techniques have facilitated large-scale tissue analysis, existing tools that automate muscle morphometry analysis are largely tailored to mammalian models, with limited applicability to teleosts. Moreover, there is a lack of effective tools for visualizing spatial organization and morphometric variability of teleost muscle fibers, a feature that is important for understanding hyperplastic muscle growth dynamics in teleosts. In this study, we show that cytoplasmic staining combined with deep learning–based cell segmentation offers a robust and accurate approach for automated muscle morphometry analysis in developing zebrafish. We also introduce a FIJI2 plugin, implemented in Jython, that streamlines both morphometric analysis and visualization. This tool accommodates shallow and deep learning–based segmentation techniques and incorporates novel quantification and visualization methods suited to teleost-specific muscle features, including mosaic hyperplasia dynamics. The plugin features an intuitive graphical user interface and is designed for flexibility, with minimal constraints regarding species, image quality, or staining protocol. Its modular architecture allows it to be used as a baseline for automated muscle morphometry analysis, while permitting integration with other tools and workflows.

## Main

Quantitative analysis of muscle morphology, or muscle morphometry, is a well-established method for investigating muscle dynamics across processes such as growth, homeostasis, disease progression, regeneration, and aging. Traditionally, this analysis has relied on manual assessment of muscle fiber size, number, and shape from histological sections—a process that is labor-intensive and time-consuming. Recent advances have introduced automated tools to streamline muscle morphometry analysis, particularly in mammalian systems such as humans and mice (Danckaert et al., 2023; Encarnacion-Rivera, Foltz, Hartzell, & Choo, 2020; Wen et al., 2018). However, these tools often depend on specific staining protocols, including precise extracellular matrix (ECM) or membrane markers (Danckaert et al., 2023; Mayeuf-Louchart et al., 2018; Wen et al., 2018). While such stains yield consistent results in mammalian tissues, their performance is frequently suboptimal in other taxa, such as teleosts—valuable but under-supported models in muscle biology. Furthermore, existing algorithms are typically designed under the assumption that muscle fibers are relatively uniform in size, a feature that aids in distinguishing true fibers from imaging artifacts (Danckaert et al., 2023). While this assumption holds true for most mammalian muscle, they fail to capture the heterogeneity characteristic of teleost muscle. Teleosts exhibit extensive hyperplastic muscle growth throughout development, resulting in a broad distribution of fiber sizes in the developing muscle (Mommsen, 2001). As a result, algorithms optimized for the structural uniformity of mammalian muscle often misclassify small, newly synthesized muscle fibers as artefacts when applied to teleost tissues. Consequently, there is a pressing need for more adaptable and species-agnostic approaches to automated muscle morphometry analysis that do not depend on narrowly defined staining protocols, sample quality, and morphological assumptions.

In this study, we demonstrate that cytoplasmic staining of muscle fibers, when combined with deep learning-based image segmentation, enables accurate and automated muscle morphometry analysis in developing zebrafish. To facilitate this, we introduce a complete, user-friendly workflow implemented as a plugin for the ImageJ2/FIJI platform (https://imagej.net/software/fiji/). The plugin provides a modular graphical user interface (GUI) that guides users through the entire process—from image segmentation to morphometric quantification and visualization. Users can select from shallow learning algorithms for faster, less resource-intensive segmentation, deep learning models for higher accuracy and throughput, or a combination of both depending on experimental needs. All approaches support custom model training using user-specific datasets, allowing for optimization across diverse sample types. Shallow models can be trained within minutes, whereas deep learning models may require several hours. Nevertheless, we show that the pre-trained Cellpose cyto3 model already performs well in segmenting muscle fibers based on cytoplasmic staining (Stringer, Wang, Michaelos, & Pachitariu, 2021), which we were able to further improve by training a zebrafish-specific model for fiber segmentation.

Following segmentation, the plugin provides tools for region-of-interest (ROI) generation and filtering, manual refinement of detected ROIs, and detailed quantification and visualization of fiber size distributions and morphometric heterogeneity. These outputs offer both visual and quantitative insights into the structural organization of teleost muscle.

We anticipate that this plugin will serve as a foundational platform for automated muscle morphometry analysis in teleosts, while retaining the flexibility needed to accommodate a wide range of species, developmental stages, and experimental designs. Its modular design also ensures future compatibility with evolving segmentation methods and image analysis tools.

## Results

### 1. Cytoplasmic staining combined with deep learning segmentation offers robust and accurate muscle morphometry analysis in developing zebrafish

Muscle morphometry analysis in teleost models is critical for studying muscle growth, aging, regeneration, and disease, yet it remains predominantly reliant on manual methods. Tools developed for mammalian systems are often incompatible with teleost tissues, largely due to their dependence on specific antibodies or staining protocols that perform suboptimally in non-mammalian species (Figure 1A, B). This limitation is especially pronounced in larval and juvenile fish, where extracellular matrix (ECM) and membrane staining is frequently weak or inconsistent (Figure 1B), rendering most existing segmentation algorithms—which rely on sharp ECM boundaries—ineffective. Even at stages when ECM staining becomes detectable, newly synthesized fibers arising from hyperplasia remain difficult to resolve based solely on membrane markers. These small fibers often occupy narrow gaps between ECM structures, resulting in low signal-to-noise ratios and irregular morphologies that pose significant challenges for membrane-based segmentation (Figure 1C).

**Figure 1.**
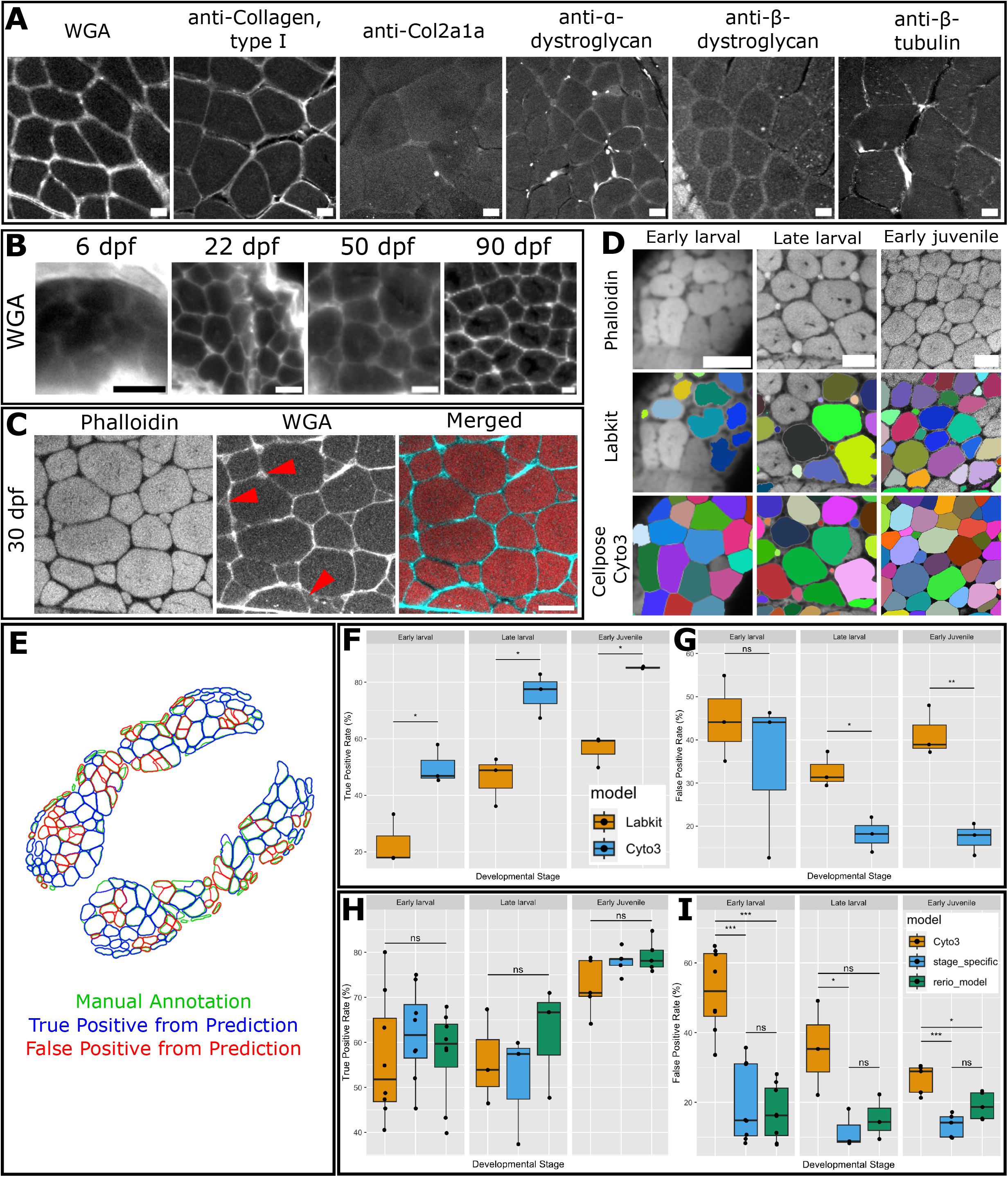
Cytoplasmic staining combined with deep learning segmentation enables robust and accurate muscle morphometry analysis in developing zebrafish. (A) ECM and membrane antibodies or stains often perform suboptimally in teleost muscle, with WGA giving sharpest signals. Images from adult zebrafish cryosections. (B) WGA signal is markedly reduced in younger zebrafish, hindering segmentation. Images from zebrafish cryosections. (C) Narrow ECM gaps between larger fibers frequently contain small, nascent fibers, but not all such spaces do. These gaps can be difficult to resolve due to low signal-to-noise ratios. Red arrows indicate ECM gaps, with and without nascent fibers. Images from vibratome sections. (D) Phalloidin consistently provides sharp cytoplasmic staining at all developmental stages, but segmentations using the shallow learning algorithm Labkit performed poorly, especially at younger stages, whereas the deep learning algorithm Cellpose (cyto3 model) was better at segmenting fibers based on cytoplasmic signal. Colored ROIs represent segmentation results. (E) To evaluate segmentation accuracy, we overlaid manually annotated ROIs (green) with predicted ROIs. Predicted ROIs that matched the annotated area by more than 65% were considered true positives (blue); others were classified as false positives (red). (F&G) Across all developmental stages, Cellpose cyto3 model consistently outperformed Labkit in correctly identifying fibers. Stats: Student’s t-test. (H&I) A custom Cellpose model trained on zebrafish muscle images (rerio model), as well as stage-specific models, further improved performance by reducing the false positive rate. However, no significant difference was observed between the rerio model and the stage-specific models. Statistics: one-way ANOVA followed by Tukey’s post hoc test. Scale bars = 20 µm in all panels.

To address these limitations, we assessed cytoplasmic staining—specifically phalloidin—as an alternative to ECM-based markers such as Wheat Germ Agglutinin (WGA). While the effectiveness of WGA declines markedly in zebrafish at early developmental stages, phalloidin consistently provides strong cytoplasmic labeling across all development stages and offers improved visualization of small, nascent fibers (Figure 1D). This makes it a more suitable marker for muscles in developing teleosts.

However, even with robust cytoplasmic labeling, segmenting closely apposed fibers remains challenging, particularly in early developmental stages. To evaluate the effectiveness of general-purpose cell segmentation algorithms in this context, we compared the performance of the shallow learning algorithm Labkit (Arzt et al., 2022) and the deep learning algorithm Cellpose (cyto3 model) using phalloidin-stained confocal images of zebrafish trunk muscle cross-sections at three developmental stages: early larval (3–7 days post-fertilization, dpf), late larval (21–30 dpf), and early juvenile (30–50 dpf). Although we trained individual Labkit models for each image to optimize performance, the algorithm frequently failed to distinguish adjacent fibers, especially in early larval tissues where fibers are tightly packed (Figure 1D). Labkit’s performance improved in juvenile samples, where ECM structures more clearly delineate individual fibers. To quantitatively assess segmentation accuracy, all fibers in examined images were manually annotated and matched with algorithmic predictions based on centroid proximity and area. A match was considered valid if the predicted and annotated regions overlapped by more than 65% (Figure 1E). Results showed that Labkit performed poorly in early larval samples due to its inability to resolve adjacent fibers, but accuracy improved in late larval and juvenile tissues (Figure 1F, G). In contrast, the pre-trained Cellpose cyto3 model—despite not being trained specifically on muscle tissue—performed markedly better without the need for retraining. It consistently outperformed Labkit in correctly identifying fibers and generated significantly fewer false positives at both late larval and early juvenile stages. Notably, Cellpose correctly segmented approximately 80% of fibers in late larval and early juvenile samples, with a false positive rate below 20%, highlighting its strong baseline performance for cytoplasmic segmentation of zebrafish muscle.

### 2. Custom, sample-specific deep learning model in Cellpose further improves fiber segmentation accuracy

Having established that the baseline cyto3 model in Cellpose can segment zebrafish muscle fibers with reasonable accuracy, we next investigated whether training a custom model based on cyto3 could further improve performance. To this end, we curated a dataset of 54 confocal images of zebrafish trunk muscle cross-sections stained with phalloidin, spanning early larval, late larval, and early juvenile developmental stages. Initial segmentations were generated using a combination of shallow and deep learning algorithms, followed by extensive manual curation to produce a ground truth dataset comprising 19,840 regions of interest (ROIs).

The dataset was randomly split into 70% for training and 30% for testing. New models were trained in Cellpose using the cyto3 model as a base. To evaluate whether a single universal model or stage-specific models would yield superior results, we trained both: a universal zebrafish model using all images (hereafter referred to as the “*rerio*” model), and three stage-specific models trained on images from each developmental stage separately. Model performance was evaluated on the held-out testing set using the same matching criteria described previously: predicted and annotated ROIs were considered true matches if their centroid proximity was acceptable and if the area overlap exceeded 65%. Surprisingly, custom models did not significantly improve the true positive rate for fiber detection across developmental stages, although we observed an upward trend in performance (Figure 1H). In contrast, custom models led to a significant reduction in false positive rates at all stages (Figure 1G), indicating that training on zebrafish-specific data improves the algorithm’s ability to reject artifacts and non-fiber regions.

Additionally, no significant differences were observed between the universal *rerio* model and the stage-specific models. However, a slight downward trend in false positive rates for stage-specific models suggests that larger datasets may grant further performance gains from stage-specific-models. Together, these findings demonstrate that training a custom zebrafish model in Cellpose improves segmentation accuracy, primarily by reducing false positive detections. Moreover, a single universal model may be sufficient for robust segmentation across a range of developmental stages, though future work may clarify whether stage-specific models provide additional benefits.

### 3. Coefficient of variation in fiber area correctly identifies growth zones and quantifies mosaic hyperplasia in zebrafish

To take advantage of our ability to perform comprehensive morphometric analysis across the entire muscle section, we aimed to quantify a defining feature of teleost muscle growth: mosaic hyperplasia. This process, characterized by the continuous addition of small, newly synthesized fibers throughout the myotome, underlies much of the post-embryonic muscle growth in teleosts (Keenan & Currie, 2019; Lee, 2010; Nguyen et al., 2017) (Figure 2B). While mosaic hyperplasia has traditionally been assessed qualitatively—typically by visually identifying small fibers deep within the myotome(Biga & Goetz, 2006; Perez, Duran, Zanella, & Dal-Pai-Silva, 2023) —there remains no standardized method for its quantification, limiting comparative analyses across developmental stages, treatments, or species. Because mosaic hyperplasia increases fiber size heterogeneity within the muscle, we developed an approach to quantify and visualize this variability by calculating the coefficient of variation (CoV) in fiber cross-sectional area across localized regions of the myotome (Figure 2Aiv & Biv). As expected from previous studies, juvenile zebrafish—where mosaic hyperplasia is active— showed elevated CoV values in the deep myotome, indicating substantial local heterogeneity in fiber size. To validate this metric, we compared CoV values in the deep myotome of early larval zebrafish (which do not undergo mosaic hyperplasia) to those in early juveniles. The analysis revealed a significant increase in CoV in juvenile muscle, consistent with hyperplasia dynamics. These findings demonstrate that local CoV provides a robust, quantitative measure of mosaic hyperplasia in zebrafish and may be broadly applicable for assessing muscle growth dynamics in other teleost species.

**Figure 2.**
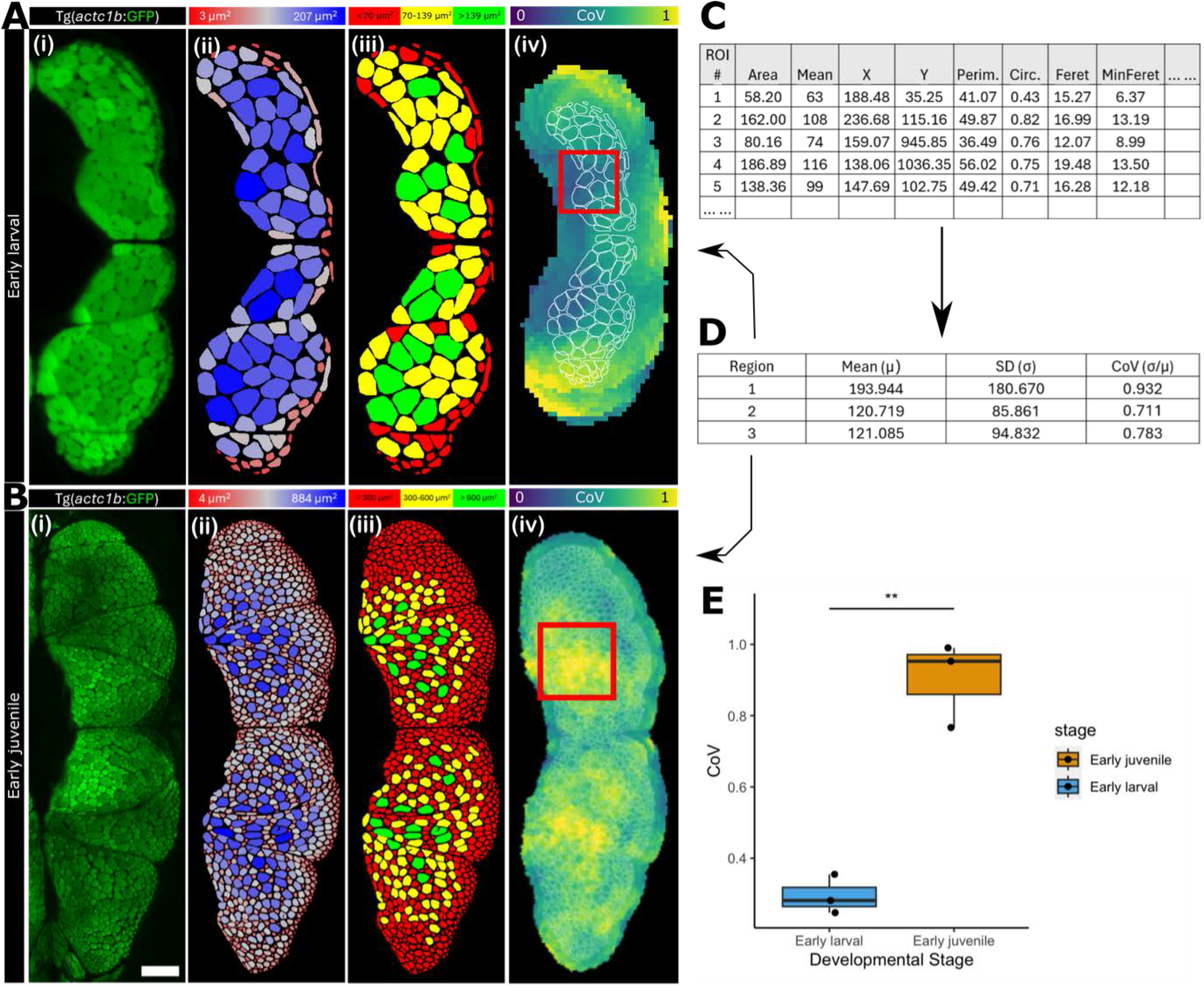
Variations heatmaps offer powerful approach to assess morphometric heterogeneity during muscle hyperplasia. (A&B) (i) Confocal images of vibratome-sectioned early larval and juvenile zebrafish showing endogenous GFP signals. (ii&iii) Identified ROIs were pseudo-colored by fiber area to generate heatmaps, facilitating visual identification of small, nascent fibers. (iv) Fiber size heterogeneity was further quantified and visualized by calculating the CoV across the myotome. (C) Heatmaps were generated based on morphometric properties of individual ROIs. (D) These morphometric parameters were used to compute local CoV across the muscle section. (E) By comparing CoV in the deep myotome of larval and juvenile fish, we demonstrate that fiber area CoV is an effective metric for quantifying mosaic hyperplasia in zebrafish.

### 4. Establishment of a start-to-finish FIJI plugin to streamline muscle morphometry analysis in teleosts

Given the complexity of options available for automated or semi-automated muscle morphometry in teleost samples—particularly the difficulties related to file format compatibility and intermediate processing steps—we developed a FIJI2 plugin designed to streamline the workflow and automate many of these intermediate tasks. In addition, we provide tools to leverage the extensive data generated through this process, including new techniques to visualize and quantify muscle morphometry, many of which have been demonstrated earlier.

As machine learning-based image segmentation continues to advance rapidly, we built the plugin’s graphical user interface (GUI) using a modular design. This modularity allows users to run each component in sequence to perform a complete muscle morphometry analysis and visualization pipeline (Figure 3A). Modules are available for both shallow learning (via Labkit) and deep learning (via Cellpose), enabling users to select the segmentation approach best suited to their samples. Similarly, dedicated modules for morphometric quantification and visualization support a range of strategies, providing users with the flexibility to adopt the most effective methods for their specific use case. Alternatively, users can integrate individual modules from this plugin into existing workflows or other software tools that may offer improved performance for their sample type.

**Figure 3.**
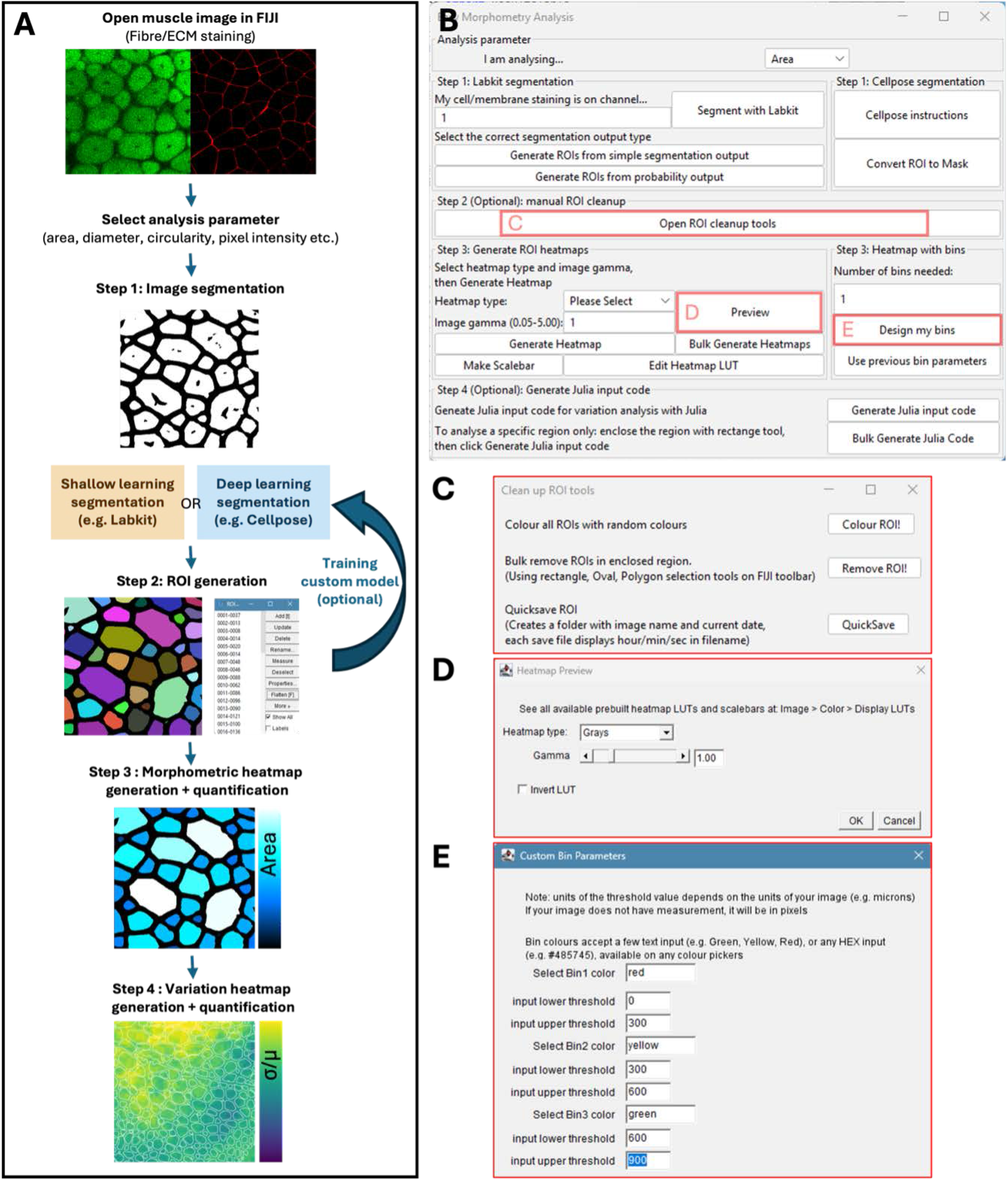
Muscle morphometry analysis workflow & UI. (A) Overview of the analysis pipeline. Our modular workflow supports both shallow- and deep-learning-based segmentation, enabling flexibility across sample types and staining strategies. We recommend deep learning algorithms for segmenting cytoplasmic staining, using our custom *rerio* model (if working with zebrafish). Alternatively, users could train their own custom models, using either shallow or deep learning algorithms to draft annotation datasets, which can then be manually corrected. (B) The plugin’s main user interface (UI) displays modules arranged in workflow order, each initiated by clicking its corresponding button. Some modules trigger additional dialog windows (indicated by red boxes) or initiate additional instructions to be followed. (C) Selecting “Open ROI cleanup tools” provides access to accessory tools for refining segmentations, including random ROI coloring, bulk deletion of ROIs circled in the image, and a quick-save button for ROIs. (D) The “Preview” function allows users to visualize heatmaps in real time under different color schemes and gamma settings, facilitating optimal visualization for their data. (E) The “Design my bins” function enables users to define custom binning criteria for ROI classification. When “Step 3: Heatmap generation” is executed, a spreadsheet containing all extracted morphometric parameters is automatically generated.

FIJI2, a free and user-friendly platform, supports a wide range of file formats through the Bio-Formats plugin (https://imagej.net/formats/bio-formats), simplifying the handling of raw microscopy data from diverse imaging systems. Additionally, FIJI2 natively supports the measurement of numerous morphometric parameters—including Feret diameter, circularity, pixel intensity, and spatial coordinates—in addition to area, all of which are incorporated into our workflow to enhance flexibility. The following section outlines the key steps of our workflow; further details are provided in the plugin’s GUI and accompanying user guide on GitHub (≤https://github.com/yansong-lu/FishROI.git≥).

### Step 1: Image segmentation

At the core of automated muscle morphometry analysis is image segmentation—a process that distinguishes true muscle fibers from background and designates them as ROIs for downstream quantification and visualization. Early segmentation strategies relied on intensity thresholding, where pixels in an image that are above a user-defined brightness value were classified as signal and the rest as noise. Although conceptually simple, this method requires uniform signal intensity and high signal-to-noise ratios—conditions rarely met in biological specimens. Recent advances in machine learning-based segmentation overcame this challenge by automatically learning a set of rules from multiple features from the image, based on annotation in the training dataset. This allows significantly improved performance, particularly in biomedical imaging and diagnostics. These algorithms generally fall into two broad categories: shallow learning and deep learning. Deep learning models offer superior accuracy but often require extensive annotated datasets, significant computational resources, and technical expertise. Furthermore, they are often highly specific to the species, tissue type, and staining used during training, limiting their utility for non-model organisms and non-standard sample preparations.

Shallow learning approaches, such as those implemented in Labkit, offer a practical alternative. They allow users to train and apply segmentation models directly on their own images using a standard laptop, often within minutes (Arzt et al., 2022). This makes shallow learning especially useful for small datasets or as means to generate initial annotations for deep learning workflows. However, as deep learning tools have become more accessible, their advantages in robustness and generalizability are increasingly compelling. As demonstrated in Figure 1F&G, the default Cellpose model (cyto3) already outperforms Labkit for segmenting cytoplasmic staining in larval and juvenile zebrafish muscle, and our custom-trained *rerio* model further improves performance. Notably, Cellpose also provides a user-friendly interface for both segmentation and model training, making it highly accessible for biologist. We therefore recommend Cellpose for processing larger datasets or for generating reusable models, particularly when working with cytoplasmic staining. For smaller projects or for creating training data, Labkit remains a useful and efficient option.

To run Labkit in our workflow, users can click “Segment with Labkit” in the plugin’s GUI. This launches Labkit in FIJI on the selected image channel. After training the classifier and saving the segmentation output, the plugin provides tools to convert the output into ROIs using the “Generate ROI from…” buttons. Note that shallow learning methods typically yield either: simple (binary) segmentations, where pixels are classified as either foreground or background, or probability maps, where pixel intensities represent the likelihood of being part of a signal. Probability maps allow users to adjust probability thresholds post-segmentation, which is especially useful when batch processing images with variable background noise. Because these segmentation types are standard outputs of shallow learning segmentation algorithms, the plugin also supports importing external segmentation maps—produced via alternative software (e.g. ilastik) —and converting them to ROIs via the same “Generate ROI…” interface (Berg et al., 2019).

If the user wishes to run the deep learning algorithm Cellpose, it is important to note that Cellpose is a standalone Python-based tool and does not operate within FIJI. To support this workflow, we provide all necessary instructions, code, and zebrafish-specific models—accessible via the “Cellpose Instructions” button in the plugin UI. Users will need to execute these commands through a command line interface (CLI) to generate the correct ROI outputs. Once segmentation is complete, the resulting outputs can be imported back into the FIJI plugin for further analysis.

Additionally, for users who wish to train new models in Cellpose, note that Cellpose does not natively accept ROI annotations in the roi.zip format generated by FIJI. To address this limitation, our plugin includes conversion tools that translate FIJI-generated ROI annotations—whether manually drawn or derived from shallow learning algorithms—into a format compatible with Cellpose training. Further details on how to train Cellpose models within our workflow can be found in the user manual available on our GitHub repository (https://github.com/yansong-lu/FishROI.git).

### Step 2: manual ROI cleanup

Regardless of the segmentation strategy used, the resulting list of ROIs is rarely perfect. As shown in Figure 1H&I, even the best-performing models fail to correctly identify some muscle fibers. Manual correction is often necessary to ensure accurate morphometric analysis. To facilitate this step, we have integrated a range of tools that allow users to visualize, manipulate, and manage their ROI sets more effectively. These include a tool that assigns random colors to each fiber, making it easier to detect instances where touching fibers have been incorrectly merged into a single ROI; a bulk ROI removal tool that deletes all ROIs within a user-defined area, avoiding the need to remove them individually; and an autosave function that allows users to periodically save ROI sets and minimize data loss in the event of software crashes—a concern especially relevant when working with large datasets or on lower-specification laptops. Once curated, the ROIs can then be quantified and visualized in the next step of the workflow.

### Step 3: Morphometric heatmap generation & quantification

In skeletal muscle, the spatial arrangement of fibers with varying morphometric properties within the myotome can reveal important biological processes. For instance, the localization of smaller fibers may indicate zones of hyperplastic growth or tissue regeneration (Figure 2A&B). To visualize such patterns, our plugin includes tools that color ROIs based on various morphometric parameters— including area, circularity, and pixel intensity—using a range of preset or custom-designed coloring schemes. Alternatively, users may assign ROIs into a defined number of bins and apply distinct colors to each bin. Regardless of the visualization strategy, a spreadsheet is automatically generated containing the full morphometric profile of each individual ROI (Figure 2B). These data can then be used for downstream analyses, such as assessing fiber size distributions to study hypertrophy or quantifying fiber numbers to evaluate hyperplasia.

### Step 4: Variation heatmap generation & quantification

Morphometric heterogeneity can provide important insights into biological processes such as mosaic hyperplasia. To support this analysis, our workflow includes the tool to quantify the CoV (CoV = standard deviation / mean) among neighboring ROIs and visualize this spatial variation across the myotome (Figure 2Aiv&Biv). These heatmaps are generated using a sliding window approach, where each pixel is colored based on the CoV between all ROIs within a defined radius (three times the mean Feret diameter) around that pixel. In addition to generating the variation heatmaps for the whole tissue, this tool will also compute the CoV within a user-defined region, allowing quantitative comparison between samples. Note that this step also requires users to run a Julia script outside of FIJI (https://julialang.org/). Our plugin generates the necessary input code by clicking the “Generate Julia Input Code” button. This code must be pasted at the end of the provided Julia script to ensure proper execution on the selected dataset. Additional instructions on installing Julia and running the script are available in our user manual on GitHub (https://github.com/yansong-lu/FishROI.git).

## Discussion

In this study, we demonstrate that cytoplasmic staining of muscle fibers is a valuable approach for accurately identifying both small and large fibers in the developing zebrafish myotome. Furthermore, we show that machine learning–based image segmentation techniques—particularly deep learning models such as those implemented in Cellpose—are well-suited for automating the segmentation of zebrafish muscle images. To accommodate the anatomical diversity and variable staining patterns from images across teleost species, we present a flexible workflow that integrates existing machine learning algorithms for muscle image segmentation, followed by novel tools for visualizing and quantifying muscle morphometry and heterogeneity. While this workflow is validated here in zebrafish as a proof of concept, the underlying machine learning strategies are expected to perform robustly across a wide range of teleost—and potentially non-teleost—species. We hope this workflow serves as a foundation for automated muscle morphometry analysis, allowing users to further expand their analyses using additional tools available within the FIJI platform.

## Methods

### Sample sources

Zebrafish husbandry was conducted according to standard procedures approved by Monash Animal Services Animal Ethics committee, Monash University (ERM41803). All zebrafish belong to the TU strain, including the wildtype and Tg(*actc1b*: GFP) transgenic line. Fish were euthanized by incubation in 1 g/L tricaine methane sulfonate. Muscles were dissected into thin blocks (∼ 10 mm) and fixed in 4% paraformaldehyde (PFA) overnight at 4 degrees. For vibratome sectioning, fixed muscle was embedded in 4% agarose and sectioned into 200 µm slices using Leica VT1200S vibratome, then stored at 4ºC in PBST until staining. For cryosectioning, fixed tissue was washed serially into 10%, 20% and 30% sucrose and embedded in Tissue-Tek® O.C.T. Compound (Sakura), followed by cryo-sectioning into 12 µm slices using Leica CM 1850 cryostats and collected on poly-lysine-coated slides, then stored at -80ºC until staining.

### Immunohistochemistry (IHC) protocol

For Tg(*actc1b*: GFP) zebrafish, vibratome sections were mounted directly into 1% low-melt agarose, followed by confocal imaging on Leica SP8 multiphoton microscope (multiphoton functionality disabled). Other zebrafish vibratome sections were incubated with 2% goat serum block (2% goat serum, 1% BSA, 1% DMSO in PBST) for 1 hour, followed by overnight incubation with Alexa Fluor 546 phalloidin (Invitrogen, A22283, 1:400 dilution) and WGA-Fluorescein (VectorLabs, FL1021, 1:300 dilution) diluted in goat serum block at 4ºC, then washed in PBST 6 times before embedding in 1% low melt agarose, followed by imaging on Leica SP8 multiphoton microscope (multiphoton functionality disabled). Cryosections were thawed at room temperature for 10 min, followed by 5 min of PFA fixation and three PBST washes. The tissue was then incubated in 2% goat serum block for 30 min, followed by incubation with WGA-Rhodamine (VectorLabs, RL1022, 1:300 dilution) for 2 hours, followed by 6 PBST washes. Sample was subsequently mounted in 80% glycerol in PBST and imaged on the Leica thunder deconvolution microscope. Additionally, we’ve also tested the following primary antibodies on cryosections after blocking, including anti-Collagen, type I (Abcam, ab23730, 1:500 dilution), anti-Col2a1a (Sigma, SAB2701915, 1:500 dilution), anti-α-dystroglycan (Sigma, 05-298, 1:500 dilution), anti-β-dystroglycan (Abcam, ab49515) and anti-β-tubulin (Abcam, ab6046, 1:500 dilution). These primary antibodies were diluted in goat serum block at 1:300 dilution and incubated overnight at 4ºC, followed by 6 PBST washes, and incubation in appropriate Alexa Fluor 488 secondary antibodies (Invitrogen, 1:300 dilution) for 2 hours, then washed again using 6 PBST washes, followed by mounting in 80% glycerol in PBST, followed by imaging on Zeiss Z1 compound microscopes.

### Labkit and Cellpose model training

Labkit models were trained manually on a 2019 MacBook Pro laptop by carefully annotating foreground and background for individual images. Model performance was monitored during training to ensure optimal segmentation quality for each image. The resulting segmentations were exported as binary masks and converted into ROIs using the “Analyze Particles” function in FIJI. For Cellpose, custom models were trained using images and corresponding ROI annotations generated in FIJI, which were converted into segmentation mask PNG files via custom scripts. Training was performed on the MASSIVE-M3 High-Performance Computing (HPC) platform using 6 CPUs, 100 GB RAM, and an NVIDIA T4 GPU (16 GB VRAM), executed through command-line interface (CLI) prompts. Image segmentation with Cellpose was also performed on the same HPC platform, using custom Python scripts.

### Model performance validation

A custom Jython script was developed to match ROIs from manual annotations with model predictions based on centroid proximity. For each matched pair, the ROI areas were compared and matches exceeding 65% overlap were classified as True Positives. The true positive rate was calculated as the number of correctly identified ROIs divided by the total number of annotated ROIs. Predicted ROIs that could not be matched to any manual annotation were classified as False Positives, and the false positive rate was calculated as the number of unmatched predicted ROIs divided by the total number of predicted ROIs. To statistically compare model performance across developmental stages, one-way ANOVA was used for comparisons among three groups, followed by post hoc Tukey’s test where appropriate. For comparisons between two groups, Student’s *t*-test was applied. All statistical analyses were performed in R, and visualizations were generated using the ggplot2 package.

### Hardware and software requirements

Our FIJI plugin and Julia script has been tested on the following operating systems: macOS Sonoma 14.2.1, Windows 10/11 Home, Ubuntu 22.04.03 LTS (64-bit) with a minimum RAM of 8 GB. However, we recommend up to 64 GB RAM for processing very large images (5-10 GB). While the processing speed of our FIJI plugin primary depends on the CPU, Labkit and Cellpose both supports GPU acceleration to speed up image segmentation. Note that Cellpose requires a GPU that supports CUDA, such as NVDIA and Intel GPU, but not AMD GPU. We have used the following software versions: FIJI2 v1.54f, Labkit 0.3.11, Julia v1.10.0. Further troubleshooting guide can be found on our GitHub page.

## Data and Plugin availability

All plugin, scripts, user manual and all models included in this manuscript is accessible on GitHub (https://github.com/yansong-lu/FishROI.git).

## Acknowledgements

We thank all aquatics technicians at Monash University FishCore for assistance in teleost husbandry. We thank Monash Micro Imaging (MMI) for its aid in microscopy. We thank Dr. Mervyn Dauer for his advice on heatmap designs. M.P. was supported by a postdoctoral research fellowship from the School of Mathematics and Statistics at the University of Melbourne. The Australian Regenerative Medicine Institute is supported by funds from the State Government of Victoria and the Australian Federal Government. The authors declare no conflicts of interest.

## Notes

### Competing Interest Statement

The authors have declared no competing interest.

